# Herbivory-triggered assemblage of sunflower rhizosphere microbiome enhances herbivore tolerance through plant–soil feedback

**DOI:** 10.64898/2026.01.28.701931

**Authors:** PM Rodríguez-Blanco, G Zitlalpopoca-Hernández, MG González Holgado, I Fernández, A Ossowicki, VJ Carrión, L Carro, A Martínez-Medina

## Abstract

**Background:** Microbial communities in the rhizosphere are key drivers of plant immunity, mediating plant responses to stress. Under specific stresses plants are capable of recruiting beneficial microorganisms into their rhizosphere with the potential to alleviate these stresses. Among these stresses, herbivorous pests remain a major agricultural challenge. Despite this, the impact of leaf herbivory on root-associated microbiomes, and how this impact can shape plant defense phenotypes are still understudied. In this study, our main objective was to determine the extent to which leaf herbivory affects the rhizosphere microbiome, and whether and how these herbivory-induced changes modulate plant defense phenotypes through plant-soil feedback. To that end, we designed a two-phase assay in which we challenged sunflower (*Helianthus annuus L.*) with *Spodoptera exigua* and later tested the effect of the microbial legacy after infestation on sunflower defense phenotype, considering resistance and tolerance as major drivers.

**Results:** We found that herbivory triggered significant changes in the bacteriome structure and dynamics, and microbiome functional profile, while effects on mycobiome were comparatively less pronounced. Under herbivory, several bacterial taxa and functional groups were enriched, the bacterial co-occurrence network was more complex and assembly processes were slightly more stochastic. Furthermore, after evaluating the plant-soil feedbacks of herbivory-induced microbiomes we observed no effect on plant resistance proxies such as herbivore growth and survival, and leaf phenolic and flavonoid content. We did observe differences on tolerance proxies, while plants grown on herbivore-challenged microbiome were overall smaller, the biomass loss to herbivory was significantly lower while the elemental nutrient content and photosynthetic pigments content was enhanced.

**Conclusions:** Our study demonstrates that insect herbivory by *S.exigua* reshapes sunflower rhizosphere microbiome and generates a soil legacy that promotes herbivory tolerance on subsequent plant generations. This highlights the broader potential of microbiome-mediated plant–soil feedbacks in shaping plant adaptation to herbivory.

## BACKGROUND

Upon herbivore attack, plants defend themselves either by reducing damage from herbivores (resistance) or by minimizing its negative fitness costs through compensatory mechanisms (tolerance) [1]. Resistance mechanisms include physical barriers like spines and chemical defenses, such as specialized metabolites that directly reduce herbivore performance [2]. In contrast, tolerance mechanisms mitigate the negative consequences of herbivory, without directly targeting the attacker. These strategies involve, among others, increased nutrient uptake and reallocation or enhanced photosynthetic rates [3, 4]. Resistance and tolerance are not mutually exclusive, and plants often deploy them in combination [5]. While anti-herbivore resistance has been extensively studied and its molecular basis is relatively well understood [2], herbivory tolerance remains comparatively understudied, despite its evolutionary relevance and its potential to mitigate yield losses under herbivory pressure [6].

Plants also provide a niche for diverse microbial communities including bacteria, fungi, protists, nematodes, and viruses, collectively constituting the plant microbiome [7]. A significant proportion of photosynthetically fixed carbon is allocated to sustaining rhizosphere microbiota, and, in return, beneficial rhizosphere microorganisms provide essential services to the host plant, such as enhanced nutrient acquisition and protection against pathogens and herbivorous pests [8, 9]. Previous studies have highlighted the relevance of the rhizosphere microbiome in plant adaptation to herbivory. Certain members of the rhizosphere community can modulate plant immunity, priming the accumulation of specialized metabolites such as flavonoids and phenolics that deter herbivores, a phenomenon referred to as microbe-induced resistance [10]. However, despite the recognized influence of rhizosphere microbes on plant resistance, their role in shaping plant tolerance to herbivory remains less understood [11].

Importantly, the rhizosphere microbiome is not static; it can be dynamically recruited and assembled throughout the plant’s life cycle. Under biotic and abiotic stress conditions, plants often alter the composition of their root exudates, likely to selectively recruit beneficial stress-tolerant microbes [12]. For example, pathogen infection causes alterations in root exudation patterns that result in the selective recruitment of resistance-inducing microbial taxa [13, 14]. Emerging evidence suggest that herbivore attack can similarly reshape exudation patterns, leading to the recruitment of potentially beneficial microbiota. Aphid and whitefly infestation, for instance, trigger changes in root exudates, associated with the enrichment of rhizosphere bacteria such as *Paenibacillus* or *Pseudomonas*, which possess insecticidal properties [15, 16]. These plant-mediated shifts can generate soil legacy effects, improving the performance and survival of subsequent plant generations via plant-soil feedback (PSF) [17, 18], with specific taxa playing central roles in this process [19–22]. Nevertheless, the mechanisms governing the assemblage and functioning of microbiomes after insect infestation remain poorly understood.

In this context, a growing yet still limited number of studies have shown that plant-mediated shifts in the rhizosphere communities can enhance plant defense against insect herbivores through PSF effects [21]. However, only few of them examined the mechanisms underlying how herbivore-conditioned microbiomes influence plant resistance and tolerance to herbivory, with outcomes often depending on specific plant-insect interactions [20, 23–25]. Overall, this emerging body of work indicates that herbivory can shape rhizosphere communities in ways that enhance resistance in subsequent plants. However, results remain fragmented, with positive or negligible effects depending on the biological system, underscoring the context dependency of these interactions [26, 27]. Moreover, the role of herbivore-induced microbiomes in shaping plant tolerance has received far less attention than their contribution to resistance [11].

In this work we hypothesized that the assembly of a herbivore-driven microbiome positively influences plant defense phenotypes against herbivory, considering both resistance and tolerance strategies. We used sunflower as the host plant and *Spodoptera exigua* as the herbivore in an agriculturally relevant system. Our main objective was to determine the extent to which leaf herbivory affects sunflower rhizosphere microbiome, and whether and how these herbivory-induced changes modulate plant defense phenotypes through PSF. We found that herbivory triggered significant changes in the bacteriome assembly and microbiome functional profile, while effects on the mycobiome were comparatively subtle. Our results further demonstrated that herbivory-induced microbiome shifts improved plant adaptation to subsequent herbivory, primarily by enhancing tolerance via PSF. Collectively, our findings underscore the capacity of microbiome-mediated plant–soil feedbacks to shape plant performance and adaptation under herbivore pressure in agronomically relevant systems.

## MATERIALS AND METHODS

### Soil, plants and herbivores

We collected soil from the *Dehesa,* an agroforestal ecosystem in Spain and Portugal consisting of grassland with scattered *Quercus* trees. The collection point (40.899619436374614, - 5.765967226652043) was located in an area with no anthropogenic activity in the last 30 years (**Supplementary Figure 1**). The first 10 cm of soil were removed prior to soil collection, and sieved at 5 mm *in situ* to remove large rocks, according to [21]. Part of the soil was sterilized by fractional sterilization over three consecutive days, with each cycle consisting of heating at 100-105 °C, for 30 minutes with 24h between cycles. Sunflower (*Helianthus annuus* L.) commercial cultivar LG 50.510 seeds were scarified with 20% (V/V) sulfuric acid, rinsed with water and then surface-sterilized with 10% sodium hypochlorite and rinsed again with water [28]. Seed germination was performed using sterile vermiculite at 24 °C, 60% humidity for 7 days and transplanted to pots at the start of the experiment.

*Spodoptera exigua* (Hübner) eggs were kindly provided by Prof. Salvador Herrero (Universitat de Valencia, Spain). Larvae were reared in growth chambers at 24 °C, 50% humidity and maintained with artificial diet according to [29] until infestation at instar L2. The artificial diet consisted of the following ingredients: 80 g cornflour, 25 g yeast extract, 25 g wheat germ, 1 g ascorbic acid, 0.8 g methyl-4-hydroxybenzoate, 0.05 g streptomycin, 8 g agar, and 500 ml distilled water.

### Experimental design: conditioning phase

Plastic pots (1.1 L) were filled with potting mix consisting of 1:1 (v:v) collected soil : sterile sand. Seven-days-old sunflower seedlings were transplanted into individual pots (one seedling per pot). A set of filled pots was also left without plants, and considered as bulk soil treatment (**Figure 1**). Pots were then placed in a completely randomized design in a glasshouse compartment under the following conditions 24 ± 2°C, 12-h light : 12-h dark photoperiod. Four weeks after transplanting, sunflower plants were assigned to each of two treatments: non-challenged plants, or herbivore-challenged plants. Plants in the herbivore-challenged cohort (Sun+H treatment) were infested with three 2^nd^-instar larvae, in the whole plant in V5-V6 stage [30]. Plants were covered with nylon mesh bags, and larvae were fed *ad libitum* for one week. Non-challenged plants (Sun treatment) were similarly covered with mesh bags. A total of 15 biological replicates (plants) were established for both Sun+H and Sun treatments, and 6 replicates were used for bulk soil treatment. One week post-infestation rhizosphere soil samples were collected by root-shaking, sieved at 2 mm, pooled in groups of three plants per pool and frozen at -80°C for subsequent amplicon sequencing and metagenomic analysis [31]. The remaining soil in each pot was collected and stored at 4°C for later use as soil microbiome inocula in the feedback phase. Soil from unplanted pots (bulk soil) was also collected, homogenized and sampled for amplicon sequencing and metagenomics, and stored under the same conditions.

**Figure 1:**
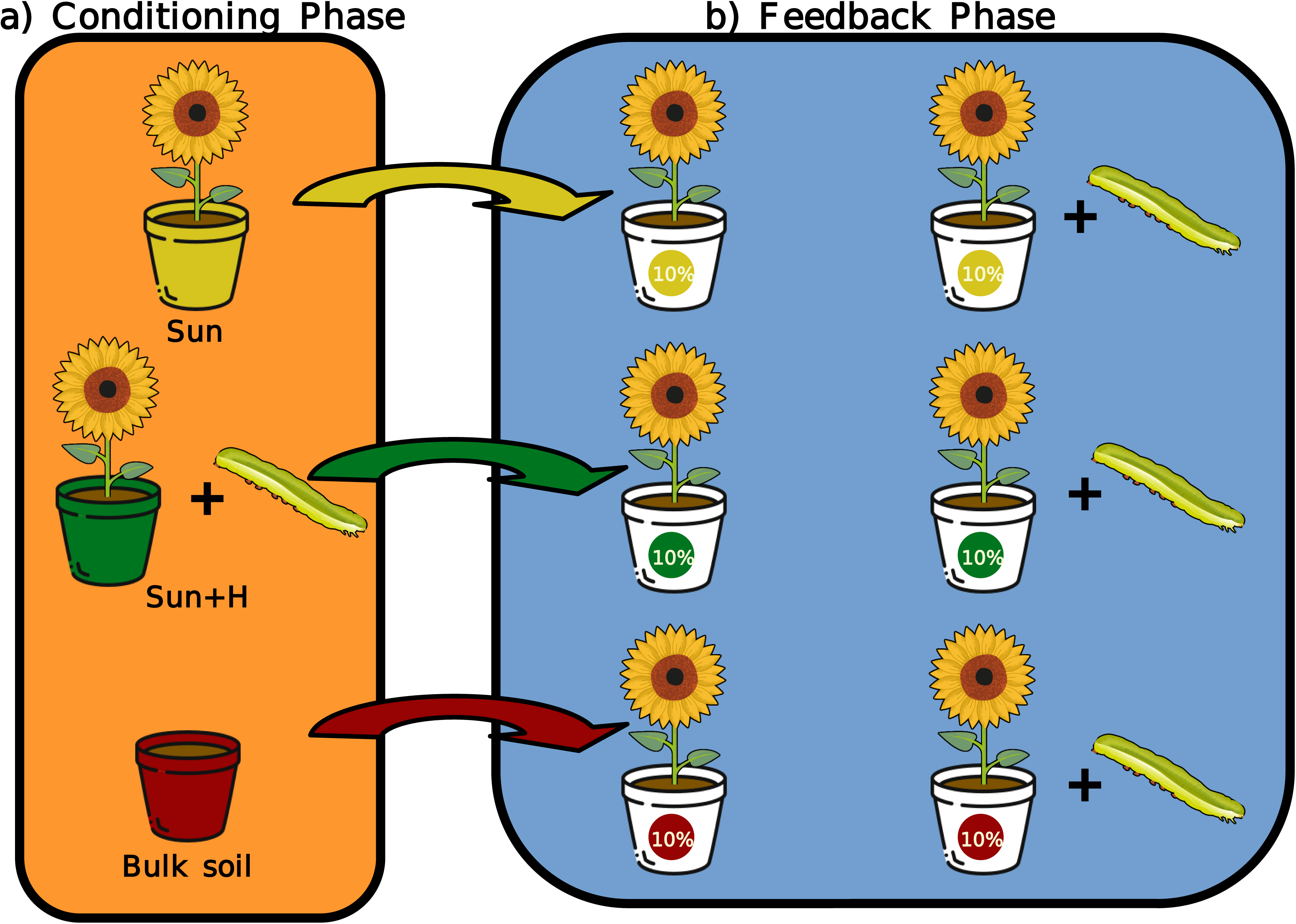
Overview of the experimental design. In the microbiome conditioning phase (a), sunflower plants were grown for four weeks, and then challenged or not with *Spodoptera exigua* for one week. This phase generated two microbial inocula associated to sunflower: Sun (from non-herbviore plants) in gold, and Sun+H (from herbivore challenged plants) in green. 15 plant replicates were grown for each, generating 15 microbiome replicates. Additionally, bulk soil from unplanted pots was used for bulk soil treatment, in red. Rhizospheric soil was sampled for microbiome analysis, including bacterial and fungal community analysis with amplicon sequencing, as well as functional profiling with metagenomics. Soils were used to inoculate sunflower plants in the feedback phase (b). In the feedback phase, new sunflower plants were grown in soils inoculated with the different treatments (Sun, Sun+H and bulk soil), and challenged or not with *S. exigua*. In the Sun and Sun+H treatments each of the 15 microbiome replicates were inoculated in one pot for each cohort (challenged and non-challenged). Effect of the different inocula on plant resistance and tolerance to herbivory traits was assessed.

### Experimental design: feedback phase

A new bioassay was setup in the glasshouse, using the three microbiome inocula produced in the conditioning phase, i.e.: microbiome inocula from Sun, Sun+H, and bulk soil treatments (**Figure 1**). Each microbiome replicate obtained during the conditioning phase was kept separated and used to inoculate two independent pots in the feedback phase, resulting in paired (“twin”) plants receiving the same microbiome inoculum. This design ensured that two plants were grown under identical microbiome conditions, allowing us to evaluate plant responses while maintaining microbiome replicate structure. Pots were filled with a sterile potting mix consisting of a 1:1 (v:v) mixture of sterilized collected soil and sterile sand. Each pot was inoculated with 10% (v:v) of the corresponding microbiome inoculum [21], and seven-days-old sunflower seedlings were transplanted into the individual pots (one seedling per pot). “Twin” plant pairs were arranged in a completely randomized design in a glasshouse compartment under controlled conditions (24 ± 2°C, 12-h light: 12-h dark photoperiod). Four weeks after transplanting, sunflower plants at the V5-V6 stage were assigned to one of two treatments: non-challenged or herbivore-challenged. Specifically, within each pair of “twin” plants, one plant was assigned to the herbivore treatment, while the other served as a non-challenged control. Herbivore-challenged plants were infested with three 2nd-instar larvae on the whole plant. Plants were covered with nylon mesh bags, and larvae were allowed to feed *ad libitum* for seven days. Non-challenged plants were similarly covered with mesh bags. A total of 15 replicates were used for each treatment, including plants inoculated with bulk soil microbiome. After 24h of herbivory the third leaf (counted from below) was sampled and stored for metabolite assessment. The weight gain and mortality of *S. exigua* larvae was recorded periodically during one week. Seven days after herbivory (5 weeks after transplanting), plant material was harvested, weighted and stored for nutrient content determination.

### Sample pooling

Samples were pooled into groups of three for microbiome, metabolite and nutrient content analyses. Rhizosphere soil was pooled prior to DNA extraction. Leaves for metabolite determination were ground with liquid nitrogen, homogenized and lyophilized together. Similarly, dry shoots for nutrient analyses were ground together. In all cases, only material from the same set of three plants was pooled together.

### Soil DNA extraction and library preparation

DNA from 350 mg from each of the 13 rhizosphere and bulk soil pooled samples was extracted using the DNeasy® PowerSoil® Pro Kit (Qiagen) according to the manufacturer’s instructions. Concentration and quality of samples was assessed using NanoDrop® spectrophotometry (Thermo Fisher Scientific). Library preparation and sequencing were outsourced to Novogene UK (Cambridge, UK). For amplicon sequencing analyses, the fungal ITS1 region was amplified using the primers ITS5-1737F and ITS2-2043R [32], while the bacterial V3-V4 region was amplified with the 341F and 806R primers for 16S rRNA gene [33]. Microbial DNA was sequenced with Illumina NovaSeq 250 bp paired-end to a sequencing depth of > 200,000 reads per sample. For shotgun metagenomic analysis, samples were sequenced on Illumina HiSeq PE150, 150 bp paired-end with 12 GB of sequencing depth per sample.

### Amplicon data analysis

16S rRNA gene and ITS amplicon sequencing data analysis was performed in R version 4.4.3 [34] and Rstudio. Quality Control and denoising into Amplicon Sequence Variants (ASVs) were performed with *dada2* v1.32.0 package [35]. For taxonomic assignment we utilized the SILVA v138.1 database [36] for bacteria and the UNITE v9.0 database [37] for fungi. *Alpha* diversity indexes (Observed and Shannon) were calculated from raw data. For the *beta*-diversity analysis we followed a compositional approach [38]. Raw data was taxonomically aggregated to family level and additive log-ratio (alr) transformed, after which euclidean distances between samples were calculated, differences in community structure visualized with Principal Component Analysis (PCA) and evaluated with euclidean pairwise PERMANOVA. To compare the differential abundance of taxa in each microbiome, a Linear Discriminant Analysis Effect Size (LEfSe) analysis was conducted using the *lefser* v1.14.0 [39] package in R. We conducted LEfSe analyses at family level with a *p-*value threshold of 0.1. Co-occurrence networks were constructed with the bacterial 16S rRNA gene amplicon dataset based on the ASV-level matrix utilizing the *NetCoMi* v1.1.0 package [40]. The 500 most abundant ASVs per treatment were retained for analysis. We considered a co-occurrence when the SparCC [41] correlation was >0.90 or <-0.90 and statistically significant (*p*-value < 0.05). Network visualization and parameters were calculated with Gephi v0.10.1 [42]. Nodes represent bacterial ASVs and edges positive or negative interactions (co-occurrences) between nodes.

Microbial assembly patterns were evaluated as described in [23]. Briefly, we calculated the *beta* nearest taxon index (*beta*NTI) for every pair of samples. A *beta*NTI >2 or < -2 reflects the predominance of deterministic processes, homogeneous selection (*beta*NTI < -2, lower phylogenetic turnover than the expected by a null model) or heterogeneous selection (*beta*NTI >2, higher phylogenetic turnover than the expected by a null model). *beta*NTI <2 and > -2 reflects the predominance of stochastic processes.

### Shotgun metagenomics data analysis

Shotgun metagenomic analysis was carried out following the SqueezeMeta v.1.6.5 pipeline [43]. Reads from every sample were subjected to quality control and pooled together and assembled into contigs with Megahit v1.2.9 [44] using kmers with lengths 27, 37, 47, 57, 67, 77, 87, 97, 107, 117, 127, 137 and 141. Contigs shorter than 500 base pairs were discarded. In the remaining contigs ORFs were predicted with Prodigal v2.6.3 [45], and taxonomically and functionally annotated into Orthologous Groups (OG) running DIAMOND v2.1.10 [46] against the nr NCBI database and eggNOG (evolutionary genealogy of genes: Non-supervised Orthologous Groups) database [47] respectively. Reads were mapped to contigs using Bowtie2 v2.5.4 [48] to generate the functional count table. To visualize the effect on the functional profile, count data was additive log-ratio transformed. We performed PCA and evaluated differences between groups with euclidean pairwise PERMANOVA. We further evaluated differences in function with an enrichment analysis using *DESeq2* v1.49.4 [49] and *ClusterProfiler* v4.17.0 [50] packages in R.

### Determination of plant nutrient and metabolite content

Nutrient content was assessed from pooled dry grounded shoot material according to [51]. Carbon and nitrogen content were determined with the Dumas combustion method (CHN628 Series Elemental Determinator). Calcium, Copper, Magnesium, Manganese, Phosphorous, Potassium, Sodium, Sulfur and Zinc content was evaluated using inductively coupled plasma atomic emission spectroscopy (ICP-OES Varian 720-ES).

Flavonoids, phenolics, free amino acids, chlorophyll, carotenoid, soluble sugar and starch content were assessed sequentially from the same material by spectrophotometry using the Rainbow protocol [52]. Briefly, we added 1 mL of 80% ethanol (4°C) to 10 mg of lyophilized pooled leaf samples and ground it in a bead mill homogenizer for 30 seconds. Samples were then centrifuged at 10,000g for 10 min at 4°C. 300 μL of the supernatant was diluted 1:1 in 80% ethanol for photosynthetic pigment content determination. The remaining supernatant was employed for total soluble sugars, total phenolic content, flavonoid content and free amino acid determinations. The pellets were incubated for 1 hr at 60°C with 1 mL of 30% of perchloric acid for starch quantification.

### Plant and herbivore data analysis

Data on plant biomass, herbivore biomass, plant nutrient content and plant metabolite content were analyzed with ANOVA models and a post-hoc Duncan test to assess the differences between groups. Normality and homocedasticity was examined prior to analysis. Herbivore survival analysis was performed with the *survival* v3.8.3 [53] package in R. Kaplan-Meier curves and LogRank test were used to determine differences between groups.

## RESULTS

### Leaf herbivory modulates rhizospheric bacteriome composition, complexity and assembly processes

We first assessed the impact of *S. exigua* leaf herbivory on sunflower rhizosphere bacteriome. We found a total of 3953 bacterial ASVs belonging to 192 bacterial families after quality control. The top ten most represented families overall in all samples were (in descending order): *Xanthobacteraceae, Solirubrobacteraceae, Nocardiodiaceae, 67-14, Pseudonocardiaceae, Micromonosporaceae, Bacillaceae, Streptomycetacea, Mycobacteraceae* and *Gemmatimonodaceae* **(Figure 2a**). These ten families represented over 55% of every sample. Notably, rhizosphere bacteriome from non-challenged sunflower plants (Sun) and sunflower plants challenged with herbivores (Sun+H) were specifically enriched in *Xanthobacteraceae* but depleted in *Nocardiodiaceae* when compared to bacteriome from bulk soil (ANOVA and Duncan post-hoc test; *p-*value < 0.05) **(Supplementary Table 1)**. However, no significant differences were found in the level of enrichment of the top ten most represented families between Sun and Sun+H treatments. Moreover, while higher *alpha* diversity was found in the communities of the Sun and Sun+H bacteriomes compared to bulk soil, no significant differences were observed between Sun and Sun+H. **(Figure 2b).**

**Figure 2:**
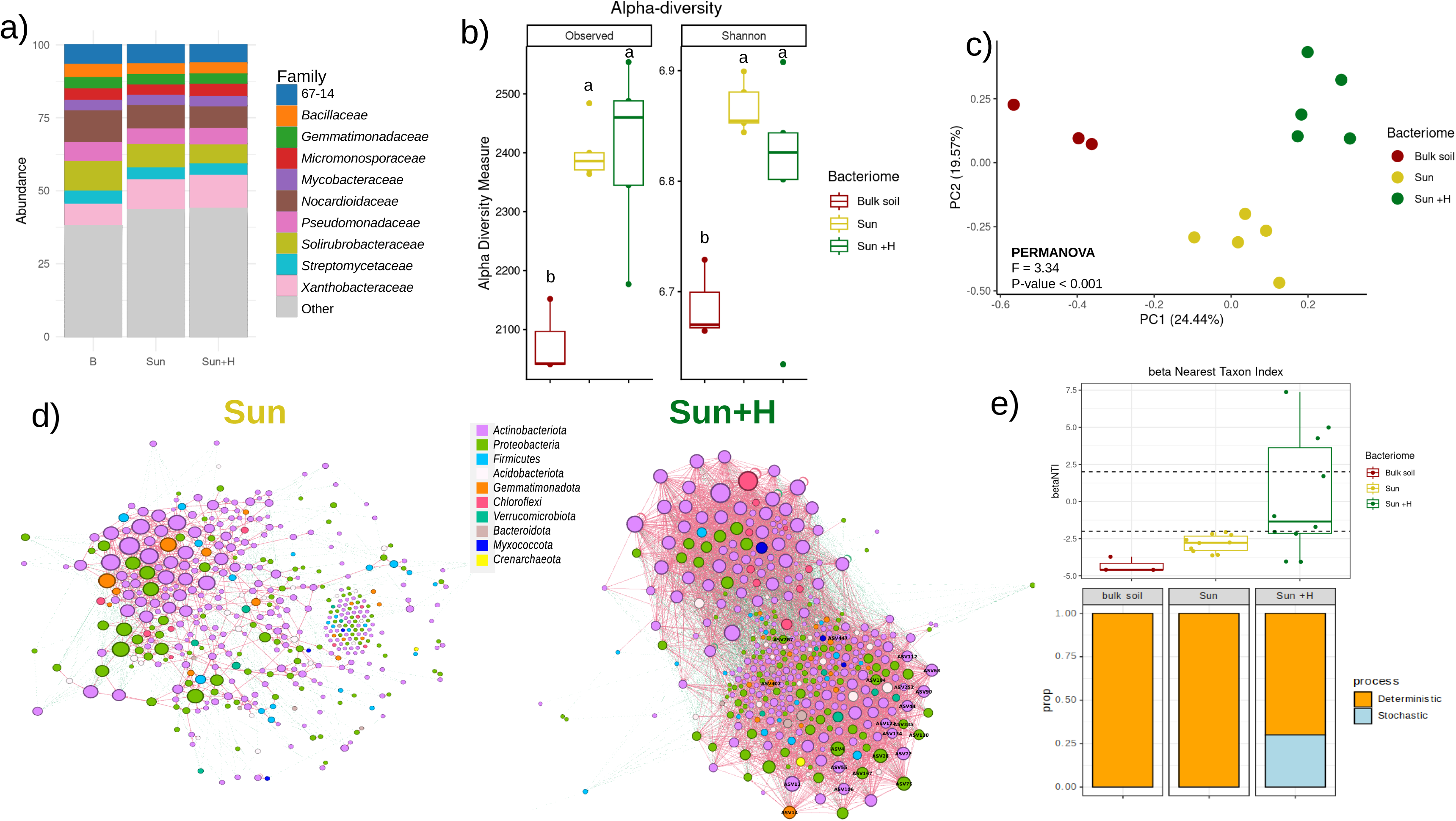
Impact of sunflower growth and *S. exigua* leaf herbivory on rhizospheric bacteriome composition, complexity and assembly. Bacteriome of bulk soil (Bulk soil), rhizosphere soil from sunflower plants non-challenged (Sun), and challenged with *S. exigua* larvae (Sun+H) were analysed. a) Stacked barplots of the top 10 most abundant bacterial families… b) Observed Richness, Chao1 index, Abundance-based Coverage Estimator, and Shannon index measures. c) Principal Component Analyisis (PCA) of the bacteriome. The community matrix is aggregated to family level and additive log-ratio (alr) transformed. X-axis and Y-axis represent the % of variability between samples explained by the first (PC1) and second (PC2) components respectively. d) Co-ocurrence network analysis of bacterial communities in Sun and Sun+H. A connection stands for SparCC correlation with magnitude >0.9 or < -0.9 and statistically significant (p< 0.05). Each node represents taxa at ASV level and is colored at phylum level. e) *Beta* nearest taxon index (*beta*NTI), defined as the difference between the phylogenetic distance between each pair of samples from a bacteriome and the distance expected from a null model (upper panel). *beta*NTI is used to estimate assembly processes in the rhizosphere, with |*beta*NTI| > 2 reflecting the dominance of deterministic processes and |*beta*NTI| < 2 reflecting the dominance of stochastic processes (lower panel). In b) different letters indicate significant differences among treatments (Duncan post-hoc test, *p*<0.05).

To compare the bacterial community structure, we then performed a PCA analysis of the additive log-ratio (alr) transformed community matrix and euclidean pairwise PERMANOVA. The analysis showed that there was an overall difference among treatments (PERMANOVA F = 3.34, R2 = 0.40, p < 0.001). Samples from bulk soil were clearly separated, mostly in PC1 (explaining over 24% of variability), from samples from Sun treatment (pairwise PERMANOVA F = 3.06, R2 = 0.34, *p* = 0.018), and from Sun +H treatment (pairwise PERMANOVA F = 3.78, R2 = 0.39, *p* = 0.018). Remarkably, there was a clear separation, mostly in PC2 (explaining almost 20% of variability) between samples from Sun and Sun +H treatments (pairwise PERMANOVA F = 1.34, R2 = 0.14, *p* = 0.048). **(Figure 2c) (Supplementary Table 2)**. Together, these results indicate that sunflower modulates bacteriome composition, and the resulting bacterial community structure differs depending on whether sunflower has been infested with an insect herbivore or not.

We further conducted a discriminant analysis with LEfSe to compare the Sun and Sun+H bacteriomes. We found that Sun+H bacteriome was enriched in the bacterial families *Xanthobacteraceae* and *Micromonosporaceae* (LDA score > |2|, *p-*value < 0.1) when compared with the Sun bacteriome. (**Supplementary Figure 2**).

We next performed network analyses to assess the impact of herbivory on the complexity of the interactions among rhizosphere bacteria. SparCC co-ocurrence networks were constructed with NetCoMi and visualized with gephi for Sun and Sun+H microbiomes (**Figure 2d**). We found that the bacterial network in Sun+H treatment (nodes = 420, edges = 8271, avg. degree = 18.927) was more complex than the network in Sun treatment (nodes = 352, edges = 1150, avg. degree = 2.632) (**Table 1).** *Actinomycetota, Pseudomonadota* and *Bacillota* were the most represented bacterial phyla in both networks. Notably, the analysis highlighted 22 hub taxa in each network. Most of these hub taxa belong to those phyla, with *Actinomycetota* being the most prevalent in Sun+H key taxa (**Supplementary Table 3)** and *Pseudomonadota* in Sun key taxa **(Supplementary Table 4)**. Hub taxa only associated with Sun+H belonged to the families TRA2-20 (order *Burkholderiales*)*, Mycobacteriaceae, Frankiaceae,* 67-14 (order S*olirubrobacterales*), *Xanthomonadaceae, Acidothermaceae, Methyloligellaceae, Propionibacteriaceae* and *Ilumatobacteraceae.* Hub taxa only in Sun belonged to *Korobacteraceae, Sphingomonadaceae, Geminicoccaceae,* JG30-KF-CM45 (phylum *Chloroflexota*), *Planococcaceae* and *Micropepsaceae*. Our findings indicate that leaf herbivory led to an increase in rhizosphere bacteriome network complexity, and highlight the potential relevance of *Actinomycetota* in the community assembly.

**Table 1:**
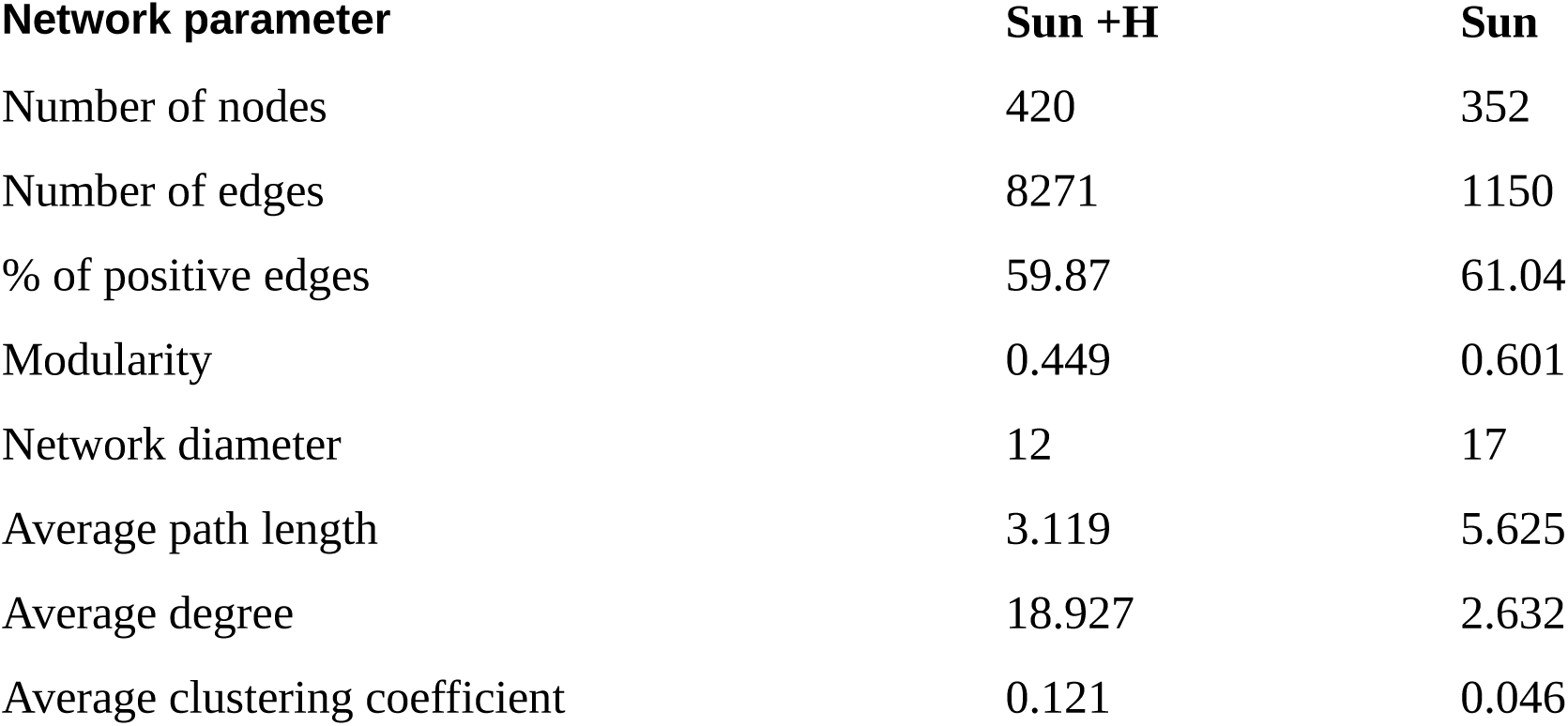
Network parameters comparison from the bacterial co-occurrence networks in Sun+H and Sun. Number of nodes refers to bacterial ASVs with at least one significant and strong (>|0.9|) SparCC correlation. Number of edges refers to the number of correlations in the network. % of positive edges are the percentage of SparCC positive correlations (>0.9) in the network. Modularity refers to the capability of the nodes to form highly connected communities. Network diameter refers to the longest distance between nodes in the network. Average path length is the average length of all edges in the network. Average degree is the average number of connections per node in the network. Average clustering coefficient refers to how nodes are embedded in their neighborhood and the degree to which they tend to cluster together.

We also explored the impact of herbivory on bacteriome assembly. We found that bacteriome community assembly was mainly driven by deterministic processes (|*beta*NTI| > 2) **(Figure 2e)**. In addition, we found that bacteriome of Sun+H treatment showed different community assembly patterns compared to bacteriome from bulk soil and Sun treatments (**Figure 2e**). Indeed, while bulk soil and Sun treatments were completely governed by deterministic processes (100% |*beta*NTI| > 2), in Sun +H part of the assembly could be attributed to stochastic processes (70% |*beta*NTI| > 2 ; 30% |*beta*NTI| < 2) **(Figure 2e).** Furthermore, in bacteriome from bulk soil and Sun treatments, deterministic processes reflect a negative phylogenetic turnover, *beta*NTI < -2, with a phylogenetic diversity lower than the expected in the null model suggesting a homogeneous selection of the bacterial communities. However, in bacteriome from Sun+H treatment, 57% of the deterministic processes reflect homogeneous selection (*beta*NTI < -2), while the reminding 43% percent reflect heterogeneous selection (*beta*NTI >2) **(Figure 2e).** Overall, our results indicated that *S. exigua* leaf herbivory modulates the rhizosphere bacteriome of sunflower, altering its composition, increasing its complexity and altering assembly processes.

### Leaf herbivory has limited effects on rhizospheric mycobiome

In analogy to bacteriome analyses, we assessed the impact of *S. exigua* leaf herbivory on rhizosphere mycobiome. We found a total of 841 fungal ASVs belonging to 151 familes after quality control. The top ten most represented families overall in all treatments were: *Herpotrichiellaceae, Mortierellaceae, Russulaceae, Aspergillaceae, Piskurozymaceae, Telephoraceae, Nectriaceae, Hypocreaceae, Trimorphomycetaceae* and Fungi_fam_Incertae_sedis **(Figure 3a)**. These ten families represent more than 66% of every sample. Remarkably, *Aspergillaceae* was depleted in Sun microbiome when compared to the bulk soil, while Sun+H and Sun were enriched in the putative family Fungi_fam_Incertae_sedis (ANOVA and Duncan post-hoc test; pvalue < 0.05) **(Supplementary Table 1)**. Mycobiome *alpha* diversity analyses did not show differences between mycobiomes from the different treatments (**Figure 3b).**

**Figure 3:**
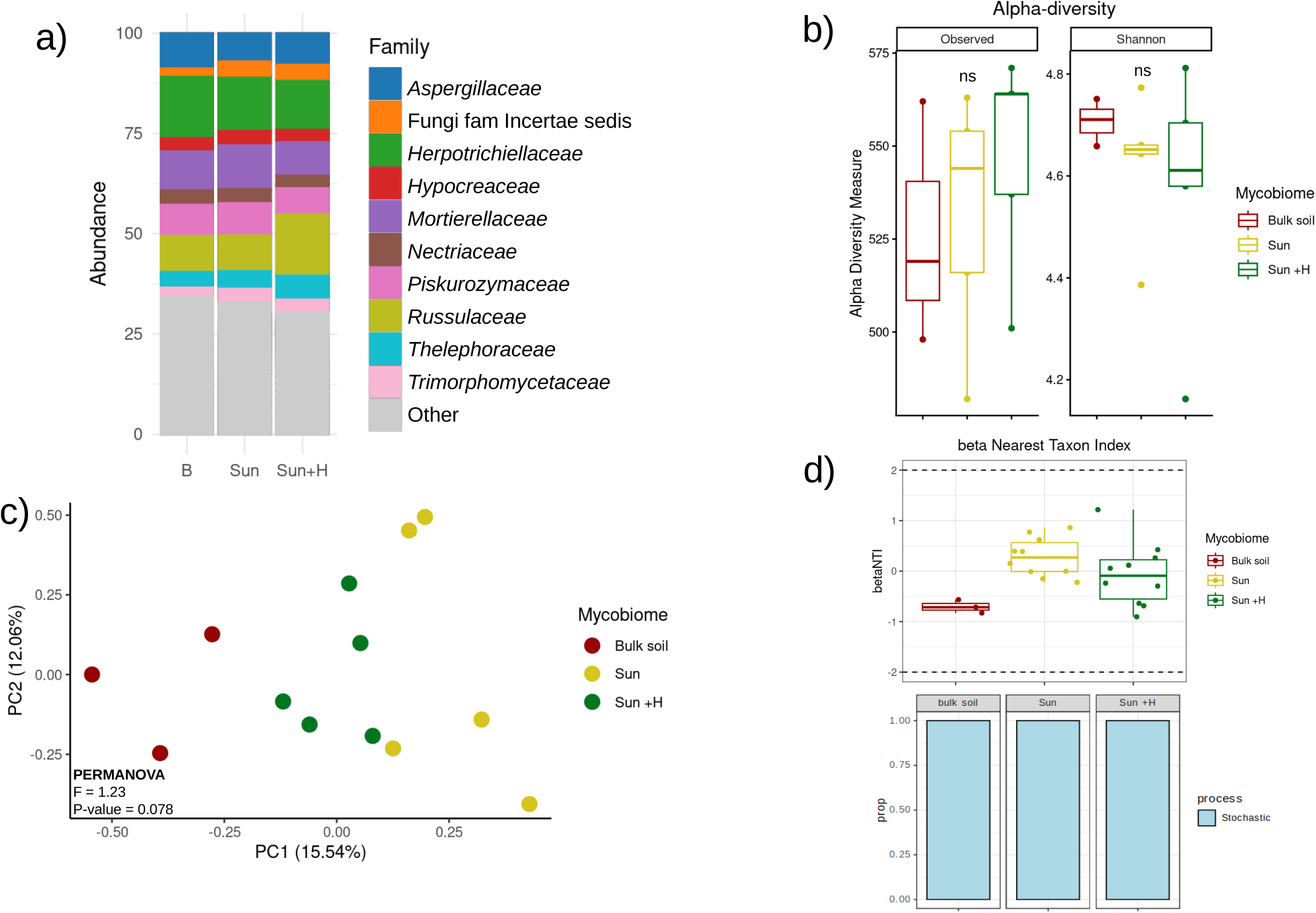
Impact of sunflower growth and *S. exigua* leaf herbivory on rhizospheric mycobiome composition and assembly. Mycobiome of bulk soil (Bulk soil), rhizosphere soil from sunflower plants non-challenged (Sun), and challenged with *S. exigua* larvae (Sun+H) were analysed. a) Stacked barplots of the top 10 most abundant fungal families. b) Observed Richness, Chao1 index, Abundance-based Coverage Estimator, and Shannon index measures in the mycobiome.. c) Principal Component Analyisis (PCA) of the mycobiome structure. The community matrix is aggregated to family level and additive log-ratio (alr) transformed. X-axis and Y-axis represent the % of variability between samples explained by the first (PC1) and second (PC2) components respectively. d) *Beta* nearest taxon index (*beta*NTI), defined as the difference between the phylogenetic distance between each pair of samples from a mycobiome and the distance expected from a null model (upper panel). *beta*NTI is used to estimate assembly processes in the rhizosphere, with |*beta*NTI| > 2 reflecting the dominance of deterministic processes and |*beta*NTI| < 2 reflecting the dominance of stochastic processes (lower panel). In b) ns: non-significant differences (Duncan post-hoc test, p<0.05).

The impact of *S. exigua* herbivory on mycobiome community structure was also analyzed with PCA of the alr transformed community matrix and euclidean pairwise PERMANOVA. The overall analysis showed marginal differences among treatments (PERMANOVA F = 1.23, R2 = 0.20, p = 0.078), separating in PC1 (that explains 15.54% of variability). Indeed, bulk soil community was marginally different from Sun (pairwise PERMANOVA F = 1.31, R2 = 0.18, p = 0.089) and Sun +H (pairwise PERMANOVA F = 1.25, R2 = 0.17, p = 0.053). However, we did not find significant differences between Sun and Sun+H (pairwise PERMANOVA F = 0.93, R2 = 0.10, p = 0.627). **(Figure 3c)**. In addition, the community assembly analysis indicated that leaf herbivory was not associated with detectable changes in the mycobiome assembly (**Figure 3d**). Fungal communities were predominantly governed by stochastic processes (|*beta*NTI| < 2) in all treatments **(Figure 3d)**. Altogether, our findings suggest that the rhizosphere mycobiome is less susceptible to alterations triggered by leaf herbivory and plant growth, compared to soil bacteriome.

### Leaf herbivory shifts the functional profile of rhizospheric microbiome

We then analysed the impact of leaf herbivory on the microbiome functions by using shotgun metagenomic sequencing. We obtained >6000000 ORFs across all samples (i.e. bulk soil, Sun and Sun+H). These were annotated into 57908 Orthologous Groups (OGs) based on the eggNOG database (evolutionary gene genealogy Non-supervised Orthologous Groups. Functional diversity analysis revealed an increase on Observed Richness index in Sun and Sun+H microbiome compared to the bulk soil, but not on Shannon index (**Figure 4a**). However, no significant differences were observed in diversity indexes between Sun and Sun+H treatments (**Figure 4a**). Interestingly, alr-transformed PCA and euclidean PERMANOVA analyses highlighted significant differences in microbiome functional structure between treatments (PERMANOVA F = 1.41, R2 = 0.22, *p* < 0.001) (**Figure 4b**). Samples from Sun and Sun+H separated from bulk soil in PC1, explaining 20.01% of the variability, while in PC2, explaining 13.53% of the variability, Sun+H samples separated from bulk soil and Sun. Specifically in the pairwise PERMANOVA, results show that Sun+H microbiome functional structure differed significantly from both bulk soil (pairwise PERMANOVA; F = 1.489; R2 = 0.199; *p* = 0.018) and Sun microbiome (pairwise PERMANOVA; F = 1.165; R2 = 0.127; *p* = 0.008) (**Supplementary Table 2**). Sun was also different from bulk soil (pairwise PERMANOVA; F =1.516; R2 = 0.202; *p* =0.008) (**Figure 4b, Supplementary Table 2**). These findings evidence a significant shift in microbiome function elicited by aboveground herbivory.

**Figure 4:**
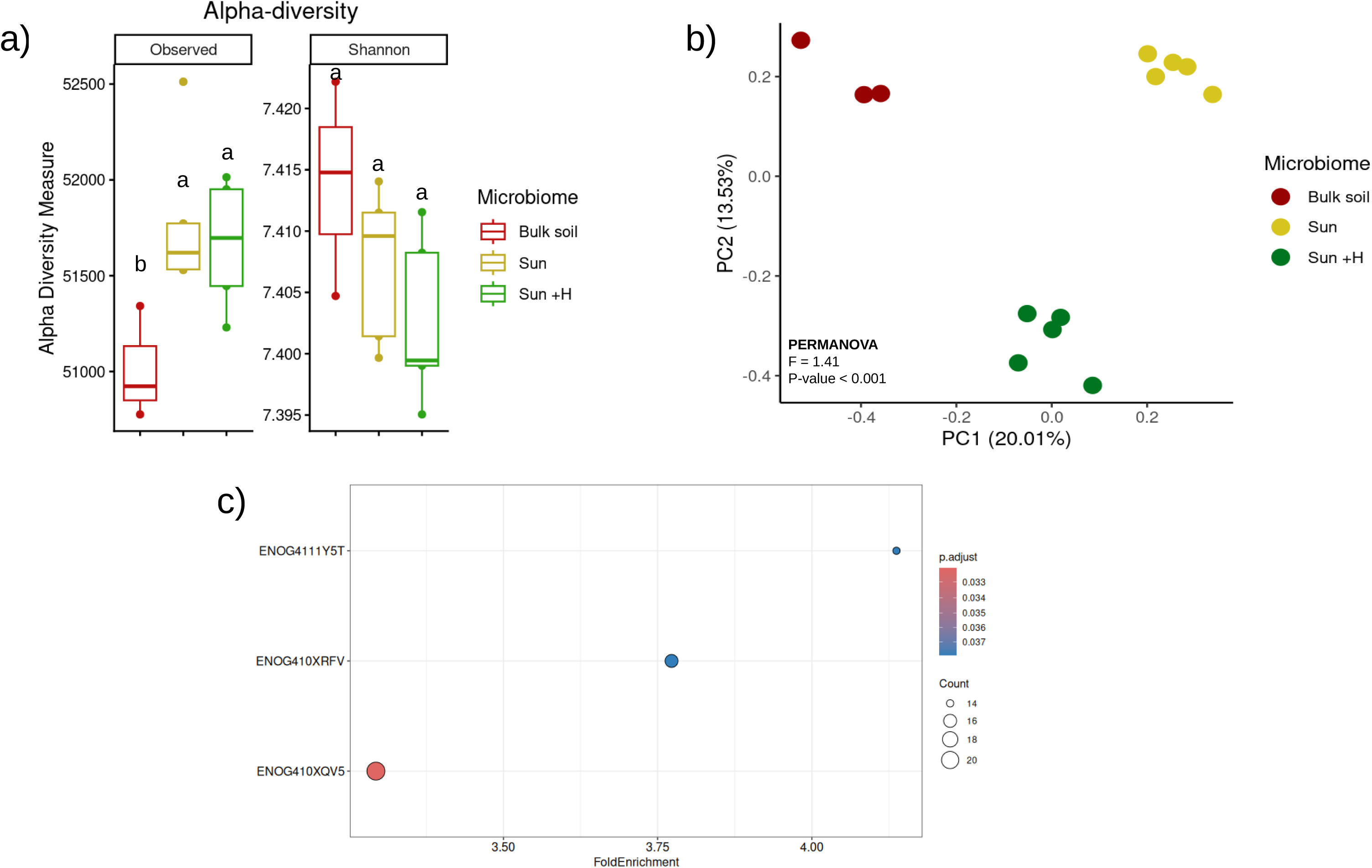
Impact of sunflower growth and *S. exigua* leaf herbivory on rhizospheric microbiome functional profile. Metagenomic analysis with samples from bulk soil (Bulk soil), rhizosphere soil from sunflower plants non-challenged (Sun), and challenged with *S. exigua* larvae (Sun+H) was performed. a) Observed Richness, Chao1 index, Abundance-based Coverage Estimator, and Shannon index measures of the functional diversity profile. Letters depict groups after Duncan post-hoc test (p<0.05). b) Principal Component Analyisis (PCA) of the microbiome functional profile annotated on eggNOG. Data is additive log-ratio (alr) transformed. X-axis and Y-axis represent the % of variability between samples explained by the first (PC1) and second (PC2) components respectively. c) Differential analysis on abundance of functional categories based on COG database. *p*-values correspond to Welch t-test. Only categories showing significant differences between Sun and Sun+H (*p*-value < 0.05) are plotted.

When performing enrichment analysis to assess specific differences among Sun and Sun+H functional profile, we found Sun+H was enriched in genes belonging to ENOG410XV5, ENOG410XRFV and ENOG4111Y5T orthologous groups from the EggNOG database (**Figure 4c**). These genes function as small conductance mechanosensitive ion channels (ENOG410XV5), triacylglycerol lipases (ENOG410XRFV) and integral membrane proteins (ENOG4111Y5T) (**Supplementary Table 5**). These results seem to indicate that herbivory alters rhizosphere microbiome function, favoring stress-responsive and metabolically active taxa adapted to herbivory treatment. Remarkably, taxonomic annotation of the enriched contigs revealed *Actinomycetota* as the main phylum containing these specific genes (ENOG410XV5, ENOG410XRFV and ENOG4111Y5T, **Supplementary Figure 3**).

### Herbivory-shaped microbiome enhances sunflower tolerance to herbivory through plant-soil feedback

We then aimed to understand whether microbiome alterations driven by leaf herbivory affect tolerance and resistance of consecutively growing plants. We performed a new greenhouse plant-soil feedback experiment. Plants were inoculated with conditioned microbiomes, including the following treatments: bulk soil, Sun and Sun+H. After growing, half of the plants were then assigned to herbivory and non-herbivory treatments. We found that in non-challenged plants, shoot biomass was reduced in plants inoculated with Sun+H conditioned microbiome, compared to plants inoculated with the bulk soil microbiome, and Sun conditioned microbiome (one-way ANOVA, F= 6.25, p = 0.0046, **Figure 5a**). We did not find significant differences in shoot biomass between plants inoculated with bulk soil microbiome and Sun conditioned microbiome (one-way ANOVA, F= 0.052, p = 0.949, **Figure 5a**).

**Figure 5:**
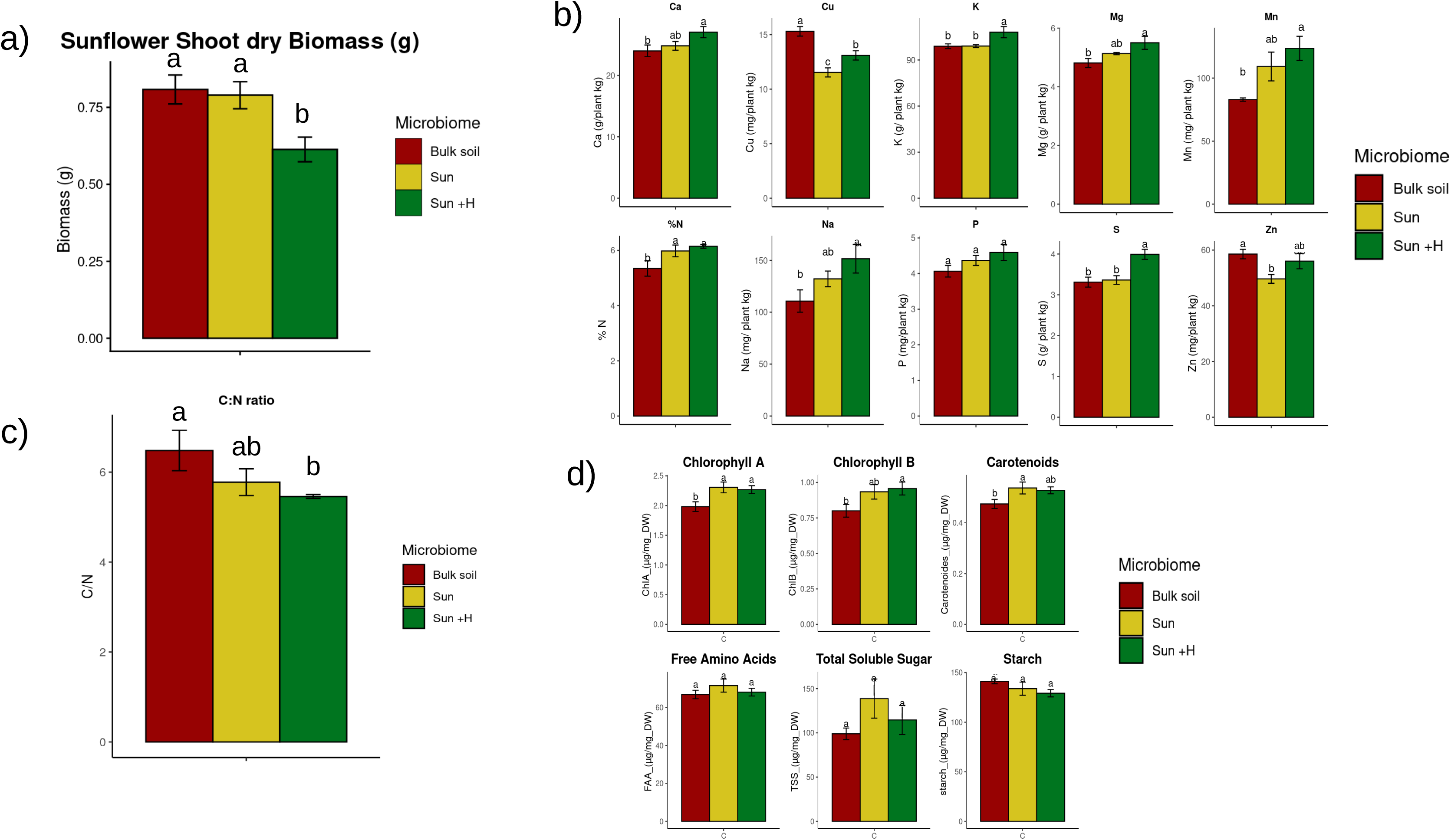
Impact of conditioned microbiomes on sunflower constitutive traits related to plant resistance and tolerance to herbivory. Plants were inoculated with the different microbiomes generated in the conditioning phase: bulk soil, Sun and Sun+H. a) Sunflower shoot biomass was assessed in non-challenged at harvest. Barplots represent mean biomass +/-Standard Error (n=15). b) Shoot content on Ca, Cu, K, Mg, Mn, N, Na, P, S, and Zn. c) Carbon-Nitrogen Shoot:Root ratio. d) Shoots content of cholorophyll A and B, carotenoids, free amino acids, soluble sugars and starch. In b) c) and d) barplots represent mean concentration and +/- SE (n=5). Different letters indicate significant differences (Duncan post-hoc test, p<0.05).

To explore the mechanistic basis of the observed effect of the Sun+H conditioned microbiome on sunflower growth, we analyzed different plant compounds relevant to herbivory tolerance, including macro and micronutrients, sugars, starch, amino-acids and chlorophyll. Overall, plants inoculated with the Sun+H microbiome showed higher levels of Ca, K, Mg, Mn, N, S and (marginally) P (one-way ANOVA, F = 2.15, p = 0.15), compared to those inoculated with bulk soil microbiome (**Figure 5b**). Inoculation with Sun conditioned microbiome also enhanced nitrogen content, compared to bulk soil microbiome (**Figure 5b)**. Remarkably Sun+H microbiome reduced Carbon:Nitrogen ratio indicating a higher relative N content, (**Figure 5c).**

Additionally, inoculation with Sun+H also enhanced the levels of chlorophyll a and b, compared to plants inoculated with bulk soil microbiome, while it had no effect on free amino-acid or sugar content (**Figure 5d**). Inoculation with Sun microbiome also enhanced chlorophyll a and carotenoid levels relative to bulk soil (**Figure 5d**). Taken together, these results indicate that herbivore-shaped rhizosphere microbiome leads to plant fortification via plant-soil feedback.

It is notable that our experimental design included “twin” plants: one infested, one non-infested, being grown with the exact same microbiome replicate. Remarkably, the impact of herbivory on shoot biomass was significantly different between the different microbiome conditioning treatments. Indeed, when comparing the shoot biomass lost due to herbivory between “twin plants” in each treatment, we found a significant lower biomass reduction in plants that had been inoculated with Sun+H conditioned microbiome, compared to plants inoculated with microbiome from bulk soil and from Sun treatment (**Figure 6a**).

**Figure 6:**
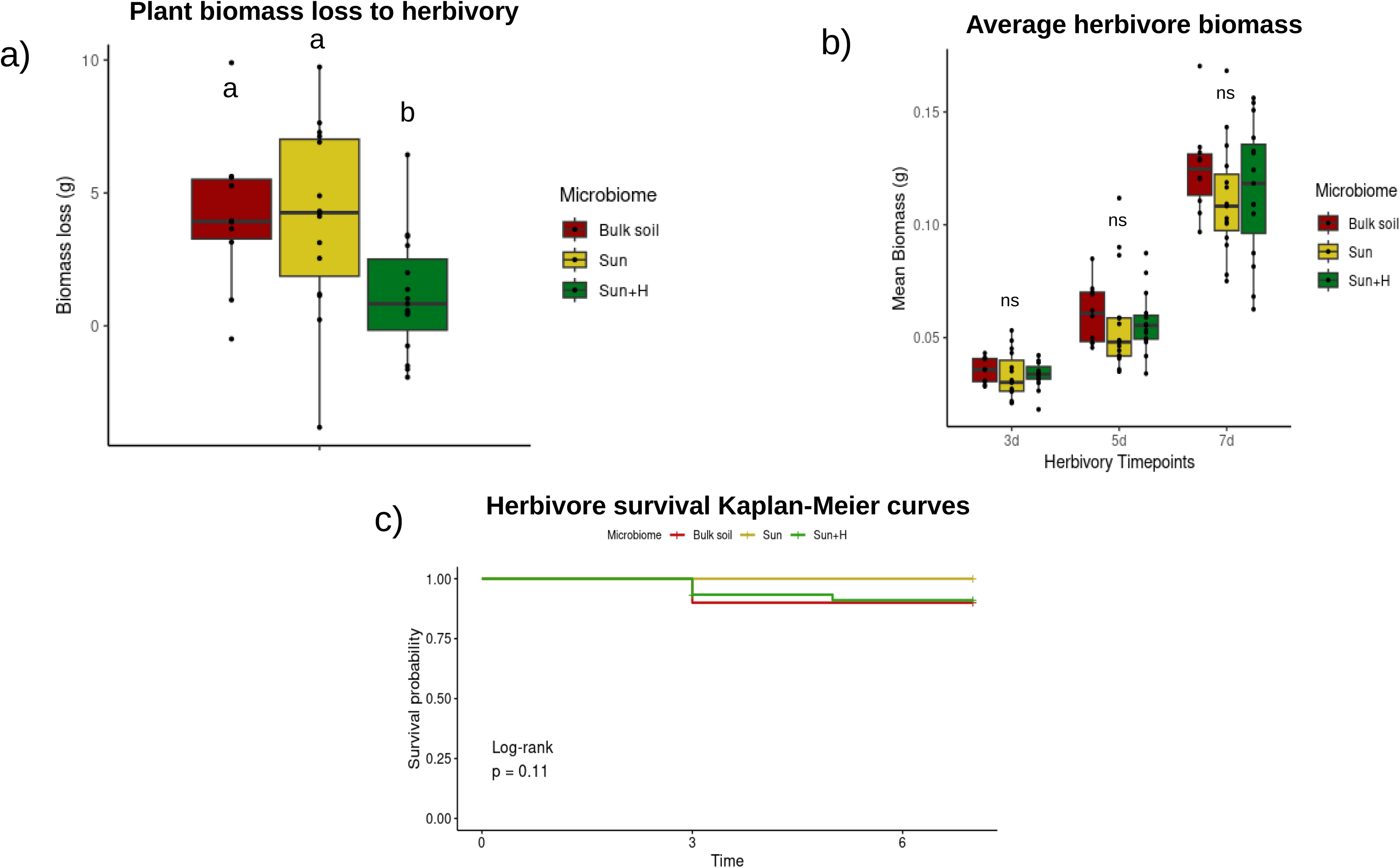
Impact of conditioned microbiomes on resistance and tolerance to herbivory . Plants were inoculated with the different microbiomes generated in the conditioning phase: bulk soil, Sun and Sun+H. a) Sunflower shoot biomass loss due to herbivory. Boxplots represent the interquartile range and median biomass loss; while whiskers represent maximum and minimum scores excluding potential outliers. b) Biomass of *S. exigua* larvae fed on plant inoculated with the different treatments, at 3, 5 and 7 days after challenge. Boxplots represent the interquartile range and median average herbivore biomass; while whiskers represent maximum and minimum scores excluding potential outliers. c) Kaplan-Meier survival curves of *S. exigua* larvae fed on plant inoculated with the different treatments, at 3, 5 and 7 days after challenge.

To explore the main drivers of this plant phenotype, we assessed the performance of the herbivores fed from plants under the different treatments. We did not find differences in herbivore biomass gain (**Figure 6b**) nor mortality (LogRank p = 0.11, **Figure 6c**) across the different treatments, indicating no significant differences in plant resistance among the different treatments. Along similar lines, we did not find differences in flavonoids and phenolic compounds across the different treatments **(Supplementary Figure 4)**. These results show that herbivory-induced changes in the rhizosphere microbiome reduces the biomass lost to herbivory in subsequently grown sunflower plants. Our findings further suggest that this effect is not driven by enhanced plant resistance, as we found no differences in herbivore performance or plant defense compounds. Instead, it seems to be driven by enhanced plant tolerance, as evidence by reduced shoot biomass loss. Moreover, the observed fortification is likely associated with this enhanced herbivory tolerance induced specifically by the Sun+H treatment.

## DISCUSSION

Insect herbivores are increasingly recognized as key drivers of shifts in rhizosphere microbial community composition and function, as highlighted by a growing number of studies [16, 22, 24]. Despite this recognition, the mechanisms governing microbiome assembly under herbivory, and the functioning of herbivore-driven microbiomes remain poorly understood, particularly in crop species. Moreover, most evidence of herbivore-induced shifts in rhizospheric microbiome comes from systems involving phloem-sucking and leaf-mining insects [16, 22, 26], whereas studies focusing on leaf-chewing insects remain scarce and often report very specific context-dependent outcomes [27, 54]. In line with our hypothesis, we found that *Spodoptera exigua* herbivory triggered significant shifts in sunflower rhizospheric microbiome, which in turn influenced the defense phenotype, specifically, tolerance strategy of subsequently grown sunflower plants.

We first found that *S. exigua* herbivory affected both the structure and the functional potential of sunflower rhizospheric microbiome. Herbivory significantly altered bacterial community structure, as revealed by multivariate analyses based on 16S rRNA gene data. In contrast, we detected no significant effects on the overall structure of the fungal community. This pattern is consistent with findings from other plant-insect systems showing that shoot herbivory can influence root-associated bacterial communities [16, 20, 24, 55], while fungal communities often show weaker or less legible responses [20, 56]. Although we do not yet have a clear explanation for why fungal communities appear more resilient to changes induced by shoot herbivory, the higher dispersal limitation of fungal communities compared to bacteria may contribute to their apparent resilience [57]. Additional studies that consider a wider range of infestation time and intensity may help determine whether chewing herbivores can meaningfully alter fungal community structure and dynamics.

The starting bacterial community showed a predominance in taxa belonging to the phylum *Actinomycetota*, an underexplored and promising phylum in insect biocontrol [58]. Among the enriched families in herbivore-driven microbiomes, *Micromonosporaceae* stands out as a potential source of insecticidal traits [59, 60]. However, while we did find enriched families, the observed shift in bacterial community structure could not be attributed to changes in specific taxa, but to a change in the whole community structure. Nevertheless, herbivory strongly altered the way the bacteriome assembles, as reflected in pronounced changes in community complexity and in the ecological processes governing its assembly. Herbivory increased the relative contribution of stochastic processes and enhanced bacterial co-occurrence network complexity. While the functional implications of the bacterial co-occurrence network complexity for plant health remain only partially understood, higher complexity has been associated to more dynamic communities and increased disease suppression capacity [61]. Interestingly, several studies examining the effects of insect herbivory on microbial co-occurrence networks reported decreased complexity [22, 23, 56], in contrast to our findings. Such differences may reflect variations in crop species, herbivore identity, herbivory intensity, and the initial soil microbial community, all of which can influence how rhizosphere communities respond to aboveground stress [20, 23]. In our system, the observed increase in network complexity may indicate that *S. exigua* herbivory promotes more dynamic and interconnected bacterial communities, potentially enhancing the functional resilience of the rhizospheric microbiome under herbivore pressure.

Beyond structural shifts, herbivory altered the functional profile of the sunflower rhizospheric microbiome, with herbivore-challenged plants exhibiting a functional profile distinct from non-challenged controls. Studies directly examining how insect herbivores alter the functional structure of the rhizospheric microbiome using shotgun metagenomic data are extremely scarce. To our knowledge, the only study reported to date did not detect herbivore-driven changes in the functional profile of rhizospheric microbiome, yet it emphasized the key role of plant-specific traits in shaping rhizosphere responses to herbivory [62]. Our results revealed an enrichment of genes encoding small conductance mechanosensitive ion channels (ENOG410XV5), triacylglycerol lipases (ENOG410XRFV), and integral membrane proteins (ENOG4111Y5T) in response to herbivory. These gene families suggest bacterial responses associated with stress sensing and adaptation in rapidly changing environments, lipid metabolism linked to plant-derived compounds, and modifications in membrane transport [63, 64]. Together, these findings indicate that aboveground chewing herbivory not only reshapes microbial community composition but also triggers functional adjustments that may enhance the capacity of the rhizospheric microbiome to sense and respond to herbivore-modified root environments.

Herbivory-induced shifts in microbial composition and function may generate legacy effects that influence the performance of subsequent plant generations [20, 25, 65]. In particular, studies on pathogen-induced changes in the rhizospheric microbiome have shown that microbial communities can modulate plant resistance, enhancing defense responses in later cohorts [19, 66]. In the case of insect herbivores, however, evidence is much more limited and fragmented, indicating strong context dependency. For example, herbivore-driven legacy effects have been shown to alter plant resistance to aphids in some studies [20, 23] and not others in similar systems [24]. In contrast, resistance to chewing insects is often unaffected [20, 23], although enhanced resistance has been reported in certain plant-insect systems [25]. In our system, microbiome shaped by S. *exigua* did not influence herbivore performance or plant chemical defenses, specifically phenolics or flavonoids. This suggests the herbivory-driven microbiome in our system does not strongly influence resistance-based defenses in sunflower.

Instead, we found a significant effect of the herbivory-induced microbiome on tolerance-related traits in subsequently grown sunflower plants. Plants grown in soils conditioned by *S. exigua* herbivory exhibited reduced biomass loss under herbivore attack due to enhanced compensatory growth. Tolerance is a critical component of plant success in both natural and agricultural systems [3], yet the role of microorganisms in tolerance remains understudied. Previous work suggests that microbial enhancement of tolerance may involve increased photosynthetic efficiency and improved plant nutritional status, among other physiological cues [4, 67]. Our results align with this idea: plants exposed to herbivore-conditioned microbiomes displayed higher nutrient content and elevated chlorophyll levels, consistent with improved photosynthetic capacity and nutrient uptake, traits known to promote compensatory growth following tissue loss. The lower C:N ratio observed in these plants further suggests enhanced nitrogen status, which could support rapid regrowth and metabolic recovery after herbivory [68]. Notably, the herbivory-induced microbiome reduced plant biomass when herbivores were absent. Although the underlying mechanisms remain unresolved, this pattern indicates a potential resource-allocation shift toward tolerance-associated metabolic pathways, generating a cost when herbivory does not occur, a scenario that is unlikely under natural field conditions where plants are rarely free from herbivore attack. Such trade-offs emphasize the context dependency of soil legacies generated by chewing herbivores and the complexity of microbiome-mediated defense phenotypes.

## CONCLUSIONS

Overall, our study demonstrates that leaf-chewing herbivory can markedly reshape sunflower rhizospheric microbiome, altering its structure, functional capacities and assembly dynamics, and generating a soil legacy capable of influencing the defense phenotype of subsequent plant generations. These shifts were particularly evident in the bacterial community and in the functional profile of the rhizosphere, pointing to the emergence of a distinct herbivore-driven root-associated microbiome. Importantly, the plant-soil feedback mediated by this microbiome enhanced tolerance, rather than resistance, indicating that microbial legacies can shape plant defense phenotypes by modulating tolerance-based strategies. Our work underlines the potential of herbivory as a driver of plant-guided microbiome selection strategies and highlights tolerance as an important part of the microbiome-induced plant response to leaf-chewing herbivores in agriculturally relevant agroecosystems.

## DECLARATIONS

### ETHICS APPROVAL AND CONSENT TO PARTICIPATE

Not applicable

### CONSENT FOR PUBLICATION

Not applicable

### AVAILABILITY OF DATA AND MATERIALS

The datasets supporting the conclusions of this article will be available in a public repository.

### AUTHORS’ CONTRIBUTIONS

PMRB and AMM planned and designed the research. PMRB, GZH, IF and MGGH performed the experiments and processed the samples. PMRB analyzed the data. AO, VC and LC provided the material resources and expertise necessary for omics data analysis. AMM secured the funding. PMRB, LC and AMM wrote the manuscript with input from all the authors.

## ACKNOWLEDGEMENTS

The authors would like to acknowledge the support of MOLECOLAB and University of Salamanca lab members that helped during the assay.

LC acknowledges the support from the ‘Escalera de Excelencia’ CLU-2025-2-04 program of the Regional Government of Castilla y León, co-funded by the Castilla y León 2021–2027 Operational Program (FEDER), Spain.

---

**Supplementary Figure 1: Soil collection point.** Experimental field situated in Muñovela, Salamanca, Spain. Part of *dehesa* agroecosystem, a semi-natural grassland-forest landscape endemic in Spain and Portugal. This particular parcel has been left with no anthropic activity for 30 years.

**Supplementary Figure 2: Discriminant analysis between the rhizosphere bacteriome of non-challenged (Sun) and challenged (Sun+H) sunflower plants.** Linear discriminant analysis Effect Size (LEfSe) determining differentially present bacterial families between Sun and Sun+H microbiomes. Differentially abundant families are considered with an LDA score > |2| and *p-*value < 0.1.

**Supplementary Figure 3: Annotated taxonomy of enriched genes in Sun+H.** Relative frequency barplot showing the phylum associated with genes in the Orthologous Groups selected in the enrichment analysis (ENOG410XV5, ENOG410XRFV and ENOG4111Y5T).

**Supplementary Figure 4: Impact of legacy microbiomes (Bulk soil, Sun and Sun+H) on flavonoid and phenolic content.** Legacy microbiomes from sunflower were selected in the Microbiome Selection Phase and its effects were tested on new sunflower plants in the Feedback Phase. Leaf total phenolic content (left). Leaf total flavonoid content. Barplots represent mean concentration and standard error (right). Letters depict groups after Duncan post-hoc test (p<0.05).

**Supplementary Table 1: Differences in abundance of the top 10 most represented bacterial and fungal families across all samples.** Letters depict groups after Duncan post-hoc test (*p*-value < 0.05).

**Supplementary Table 2: Pairwise PERMANOVAs depicting differences in the community structure and functionality of the rhizosphere.** Microbiome legacies (Sun, Sun+H and bulk soil) are considered different in the bacterial or fungal community assembly or the functional profile when *p*-value of the pairwise PERMANOVA test among them is < 0.05.

**Supplementary Table 3: Hub taxa in the bacterial co-occurrence network of sunflower plants challenged with *S. exigua*.** Hub taxa are the top most connected ASVs in the network. These are often considered keystone taxa, organisms that play an important role in the assembly of the bacterial community.

**Supplementary Table 4: Hub taxa in the co-occurrence network of sunflower plants that were not challenged with an herbivore.** Hub taxa are the top most connected ASVs in the network. These are often considered keystone taxa, organisms that play an important role in the assembly of the bacterial community.

**Supplementary Table 5: Contig table with all accessions matching the differentially abundant Orthologous Groups in the rhizosphere**. Each contig has been annotated into an Orthologous Group in the EggNOG database with its corresponding function.

