## Supplementary Figures for "Herbivory-triggered assemblage of sunflower rhizosphere microbiome enhances herbivore tolerance through plant–soil feedback"

### Slide 1
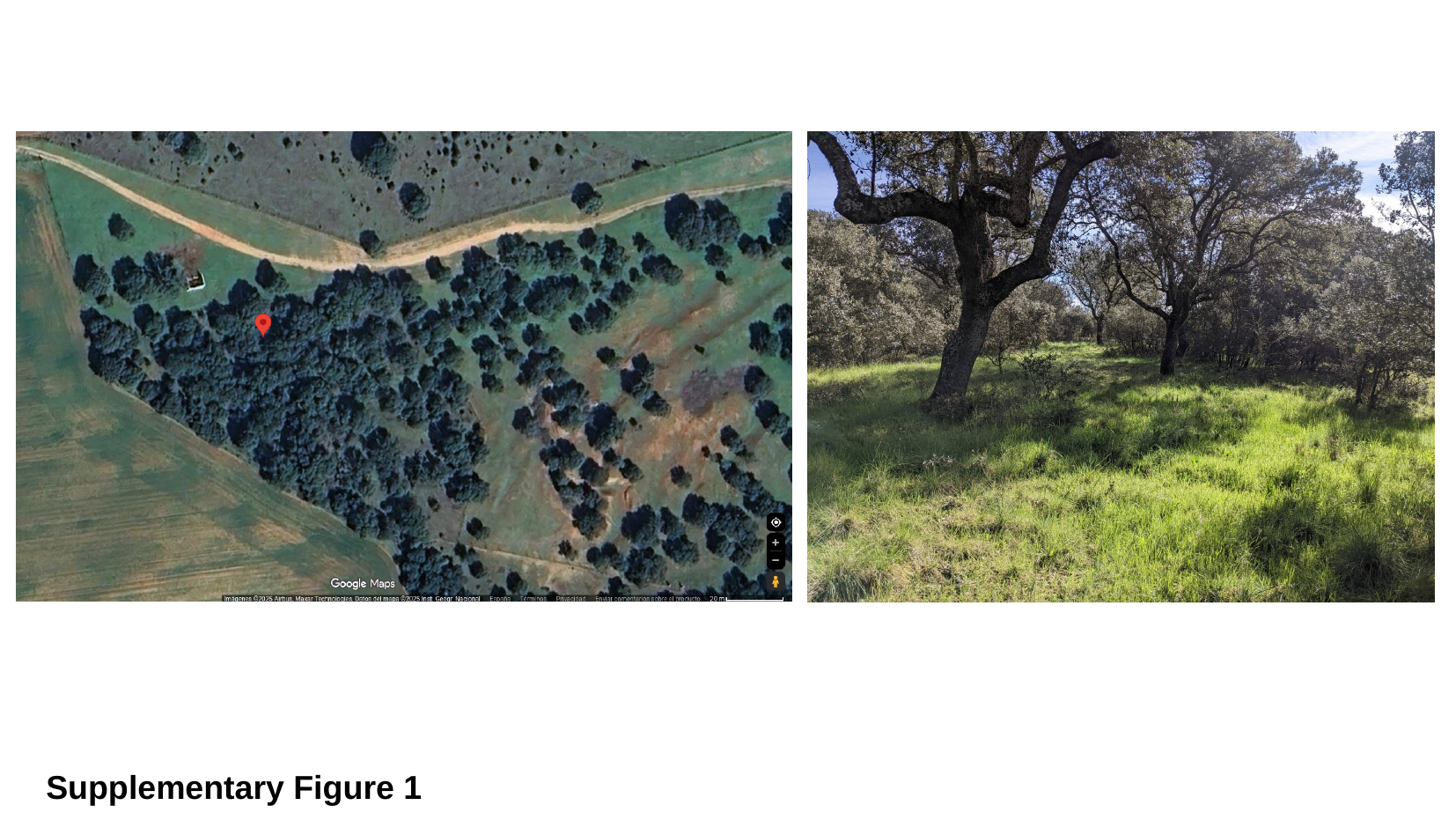

Supplementary Figure 1

### Slide 2
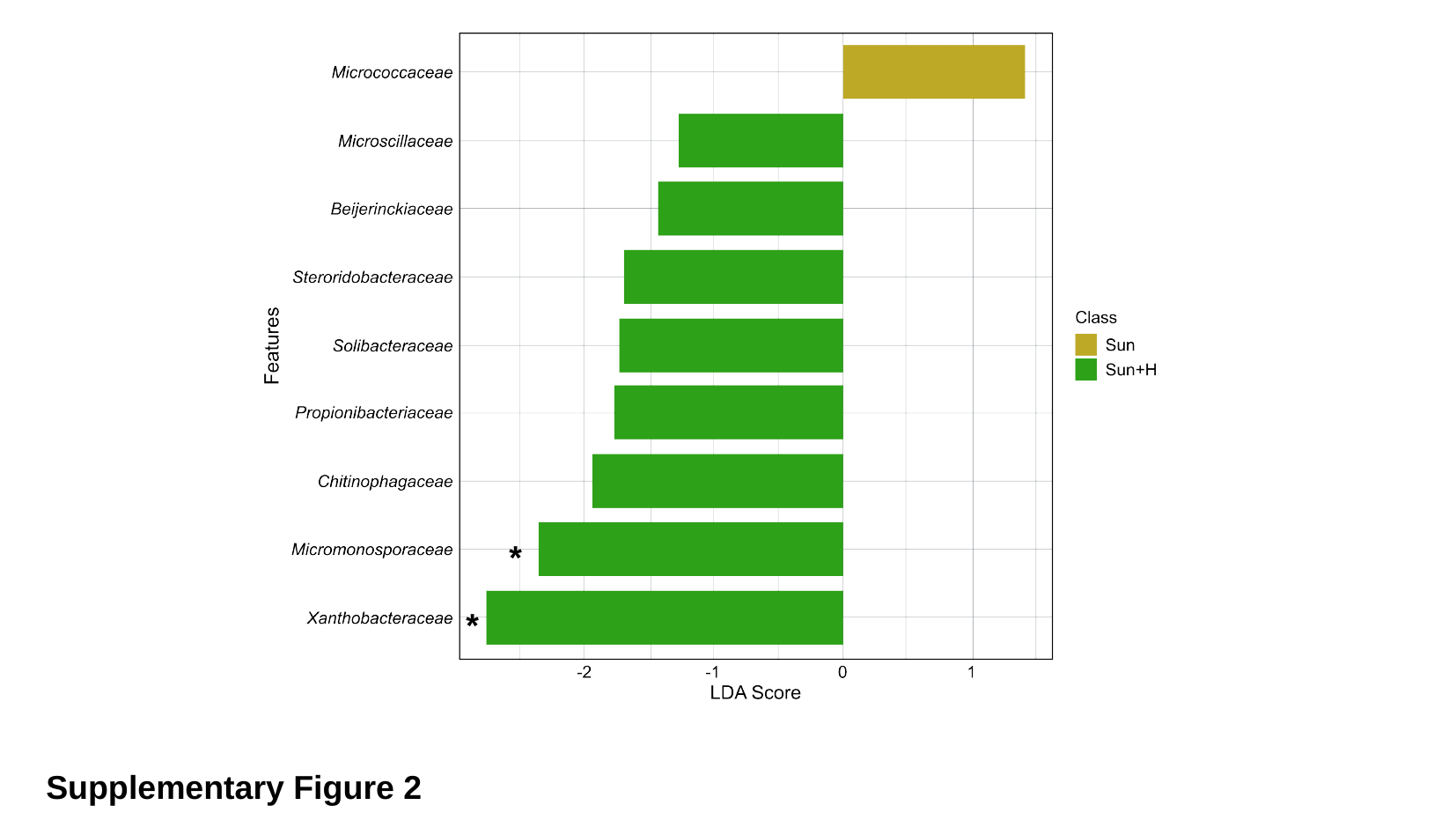

*
*
Supplementary Figure 2

### Slide 3
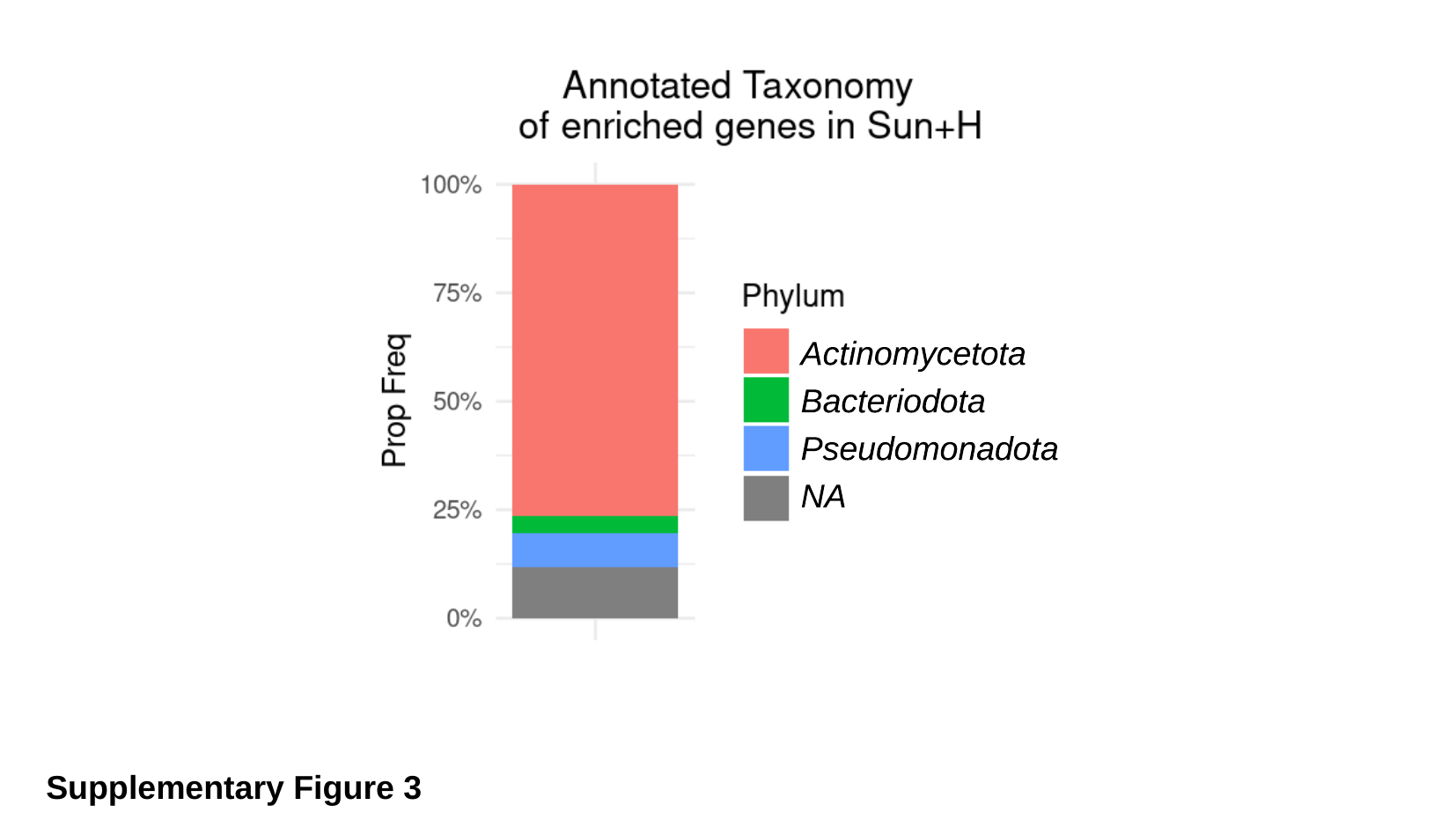

Supplementary Figure 3

### Slide 4
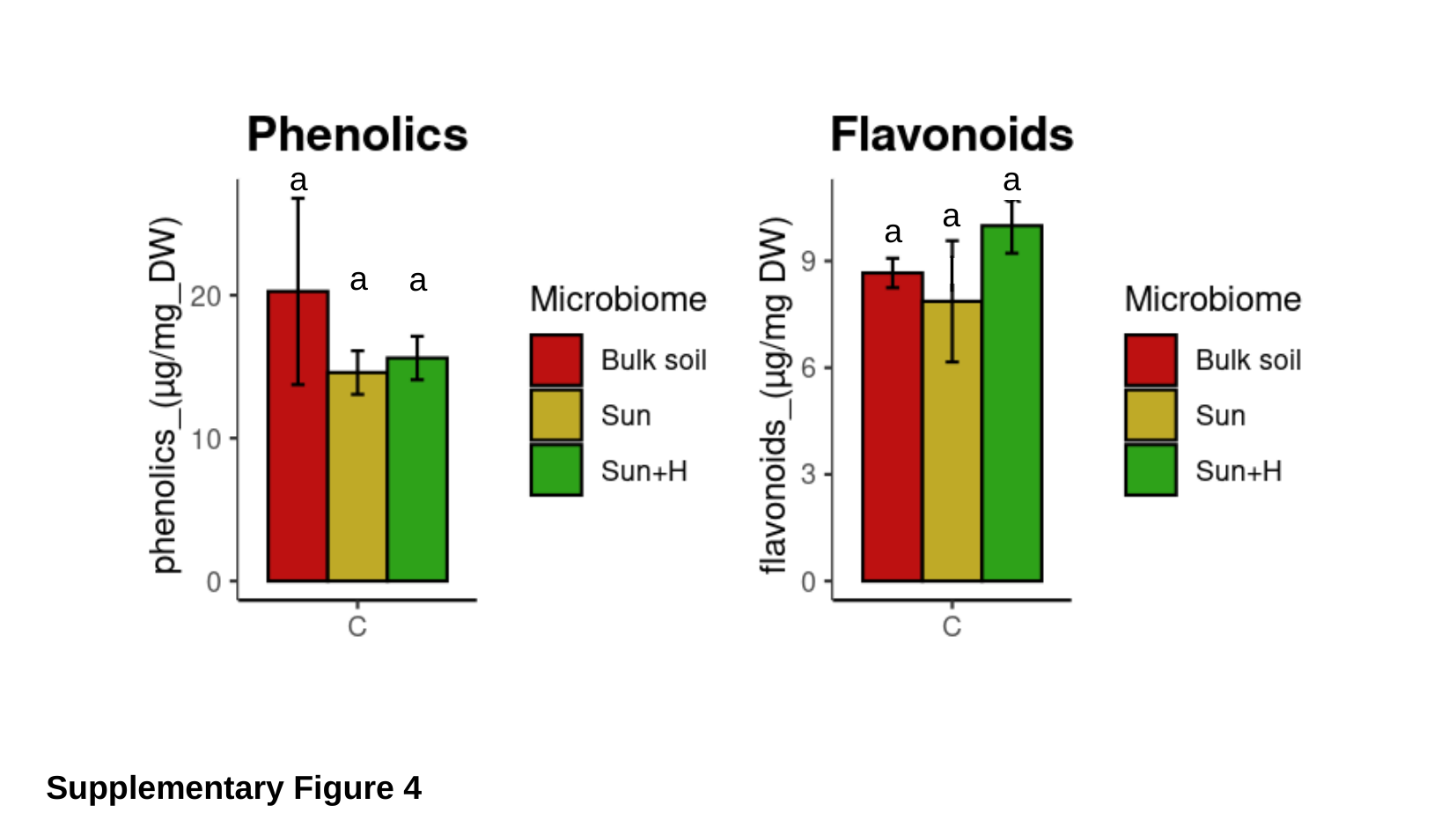

a
a
a
a
|
a
a
Supplementary Figure 4
